# A randomised exploratory investigation of the effects of Attention vs Working Memory Training on cognitive performance and everyday functioning following stroke

**DOI:** 10.1101/296723

**Authors:** Polly V Peers, Duncan E Astle, John Duncan, Fionnuala Murphy, Adam Hampshire, Tilak Das, Tom Manly

## Abstract

Difficulties with attention are common following stroke and are associated with poor outcome. Home-based online cognitive training may have to the potential to provide an efficient and effective way to improve attentional functions in such patients. Little work has been carried out to assess the efficacy of this approach in stroke patients, and the lack of studies with active control conditions and rigorous evaluations of cognitive functioning pre and post training means understanding is limited as to whether and how such interventions may be effective. Here we compare the effects of 20 days of active cognitive training using either novel Selective Attention Training (SAT) or commercial Working Memory Training (WMT) programme, versus a waitlist control group, on a wide range of attentional and working memory tasks, as well as on self-reported everyday functioning. We demonstrate separable effects of each of the active training conditions, with SAT leading to improvements in both spatial and non-spatial aspects of attention and WMT leading to improvements only on very closely related working memory tasks. Both training groups reported improvements in everyday functioning, which were associated with improvements in attentional functions, suggesting that improving attention may be of particular importance in maximising functional recovery in this patient group.

## Introduction

Stroke is the leading cause of long-term disability in the UK and other developed nations, with high costs for health and care provision [1]. It can result in persistent physical, cognitive and mood impairments. Whilst the pattern of cognitive impairment will vary according to factors such as lesion extent and location, some presentations are particularly common.

Impaired attention has been reported in up to 92% of stroke survivors in the acute stage [2] and to persist in up to 50% in the longer-term [3,4]. There are many reasons to think that attention – including our ability to detect errors and to remain focused on activities – would be critical skills in maximising functional recovery. Indeed capacity to sustain attention 2 months after stroke is a stronger predictor of motor recovery over the following two years than the level of physical impairment in the acute stage [5]. Similarly attentional functioning has also been linked to recovery of other functions such as language [6]. Moreover attentional deficits that impact spatial awareness (particularly unilateral neglect) are associated with high levels of disability, poor outcome, and increased reliance on public services [7,8].

Perhaps due to its striking presentation (including failure to eat food from half the plate, or dress one side of the body) and link to poor outcome, much of the focus on rehabilitation of attentional difficulties has focussed on trying to reduce the spatial difficulties seen in patients with unilateral neglect. Interventions specifically aimed at ameliorating the spatial bias observed in these patients, including adaptation to prism lenses [9], hemifield patching [10] and training in visual scanning [11], have had some impact. However a Cochrane review [12], concluded that there was insufficient evidence of generalised, persistent gains to currently recommend any intervention.

Given the strong evidence that attentional impairments may be key to maximising functional recovery, and that spatial interventions have not given rise to generalised persistent improvements in patients, could training non-spatial aspects of attention be beneficial? A potentially different approach to the rehabilitation of attentional impairments is informed by observations pathological spatial biases are observable in a large proportion of patients with unilateral brain lesions (not just patients with neglect) and that these spatial biases tend to arise and persist in the context of more general (not specifically *lateralised*) attentional impairments [13,14,15,16,17]. Additionally, interventions that temporarily manipulate general attentional resources during assessment of attentional functions, for example increasing alertness via stimulants or stimulation [18, 19] or reducing alertness with sleep onset [20], have been shown to phasically modulate spatial bias, suggesting rather direct interactions between these components. Despite not explicitly targeting spatial bias, therefore, it may be possible to improve spatial functions by focussing on other aspects of attention.

The distinction between lateralised and non-lateralised aspects of attention is perhaps best illustrated by computational models of normal attention, such as Bundesen’s Theory of Visual Attention (TVA)[21]. Within the framework of TVA a number of separable, but interacting components can be derived from data collected in a simple partial and whole report paradigm in which participants are requested to report the identities of very briefly reported letters of a target colour (for example black) whilst ignoring letters that may simultaneously appear in a distracting colour (say white). Some components such as visual capacity (K, how many letters can be taken in ‘at a glance’) and attentional selection (α, the degree to which the target color can be used to exclude the influence of non-target letters) are essentially non-spatial in nature. In addition the paradigm allows for computation of a distinct component of ‘spatial bias’ (a systematic bias towards/away from letters on one side of the display). Data from patient groups who have undertaken such assessment indeed confirm the link between spatial bias and more general attention capacity limitations [14,15,16]. An interesting and clinically highly relevant test of whether cognitive training of attention has generalised benefits is therefore whether gains are observed on measures of spatial bias despite patients being given no specific guidance in paying attention to the relatively neglected side of space.

There has been little scientific evaluation of the potential success of training specific cognitive functions following stroke (see [22,23] for exceptions). Westerberg et al., [23], for example, report positive findings from working memory training (WMT) suggesting improvements following training that appear to generalise to untrained tasks. The absence of an active control group, however, makes it impossible to rule out the possibility that these effects reflect the general benefit of being involved in any intervention, or participants’ expectations. Johansson, & Tornmalm [24], in contrast, detected improvements only on trained tasks. Here, in a proof-of-concept study, we ask whether computerised training, which focusses on improving attentional functions can produce specific, measurable changes in cognitive functioning and reduce disability in everyday life.

To this end we developed a novel Selective Attention Training (SAT) battery, consisting of five tasks developed to shape participants’ ability to rapidly attentionally sift through onscreen stimuli for goal-relevant information. We intended to compare this with another, well established, cognitively demanding WMT battery, Cogmed^**™**^. Working memory can be conceived as the operation of two specific, capacity/time limited, information stores (verbal and visuo-spatial) and a more general ‘central executive’ component required in many attentionally demanding activities [25]. In as much as WMT enhances capacity or efficiency of this central component, it provides a good comparison to our attentional training, with the potential to lead to improvements in attention though structurally dissimilar tasks to those developed in our SAT. WMT has been studied extensively, mainly in developmental populations. Some studies show gains which may stem from changes within the attentional control system [26,27] whilst others show that these improvements extend only to tasks that are similarly structured to those practised during training [28], suggesting that, in children at least, task-specific strategies, rather than generalised attentional improvements, may account for the behavioural gains made.

Whilst generalisation remains highly questionable within the developmental literature [29,30] there are grounds to believe that the case of stroke patients may be very different. Importantly, whilst school-aged children are exposed to hours of structured mental stimulation and feedback in the class-room each day, stroke patients in the community do not receive such stimulation, or feedback, which may be crucial to learning or relearning cognitive skills.

Taking the lead from the Cogmed WMT battery, we produced our SAT to share many of the features that have shown promise in WMT. Both forms of training are adaptive (becoming progressively more difficult as performance improves, and easier if performance is poor), allowing patients to progress at their own rate. Improvements in performance are rewarded with points, melodic flourishes or spoken feedback. Both forms of training employ varied, relatively brief tasks, using colourful displays, and provide trial-by-trial feedback to assist with learning.

Cognitive training could produce general benefits that are unspecific to the training tasks (e.g. structured daily reinforced cognitive practice, sense of confidence and mastery, general expectation effects and so forth). Comparing two forms of cognitive training provides the potential to examine both specific cognitive effects and general benefits of training (assuming a wide-ranging evaluation of cognitive functioning pre and post training is carried out to determine whether differential effects of each training paradigm are observed). To this end we randomly allocated patients with likely difficulties in attention following stroke to a WMT, SAT or WL condition. An extensive range of outcome measures assessing working memory, attention, spatial bias, and self-report everyday function were completed before and after 4-weeks of daily training (or equivalent waitlist period). To increase power, WL participants were then randomised to one or other form of training, with their post-WL assessment acting as the baseline for subsequent post-training reassessment. A further benefit of having these carefully matched training regimes, in combination with a range of theoretically motivated cognitive outcome measures is that it allows us to start to explore the potential mechanisms by which any improvements in cognitive function have occurred.

Logically three potential outcomes could be predicted, which would lead to different conclusions about associated mechanisms. Firstly, neither training regime could be associated with improvements on the outcome measures compared with the WL control. This would question whether any form of training could be effective, or whether this null finding could be due to ‘dose’ or insensitivity of the outcome measures. Secondly, both forms of training could produce equivalent general benefits compared with WL suggesting common mechanisms which could potentially be due to motivational or social influences as opposed to training specific cognitive abilities. Finally each form of training could produce its own profile of cognitive improvements suggesting mechanisms that are, at least in part, specific to each form of training. For example one might expect to find the greatest effects of WMT on working memory measures and conversely the greatest effects of SAT on attention measures. Finally, given the importance of attention for outcome in stroke, we have included a measure of everyday function to examine whether specific improvements in cognitive function influence everyday functioning.

**Figure 1.**
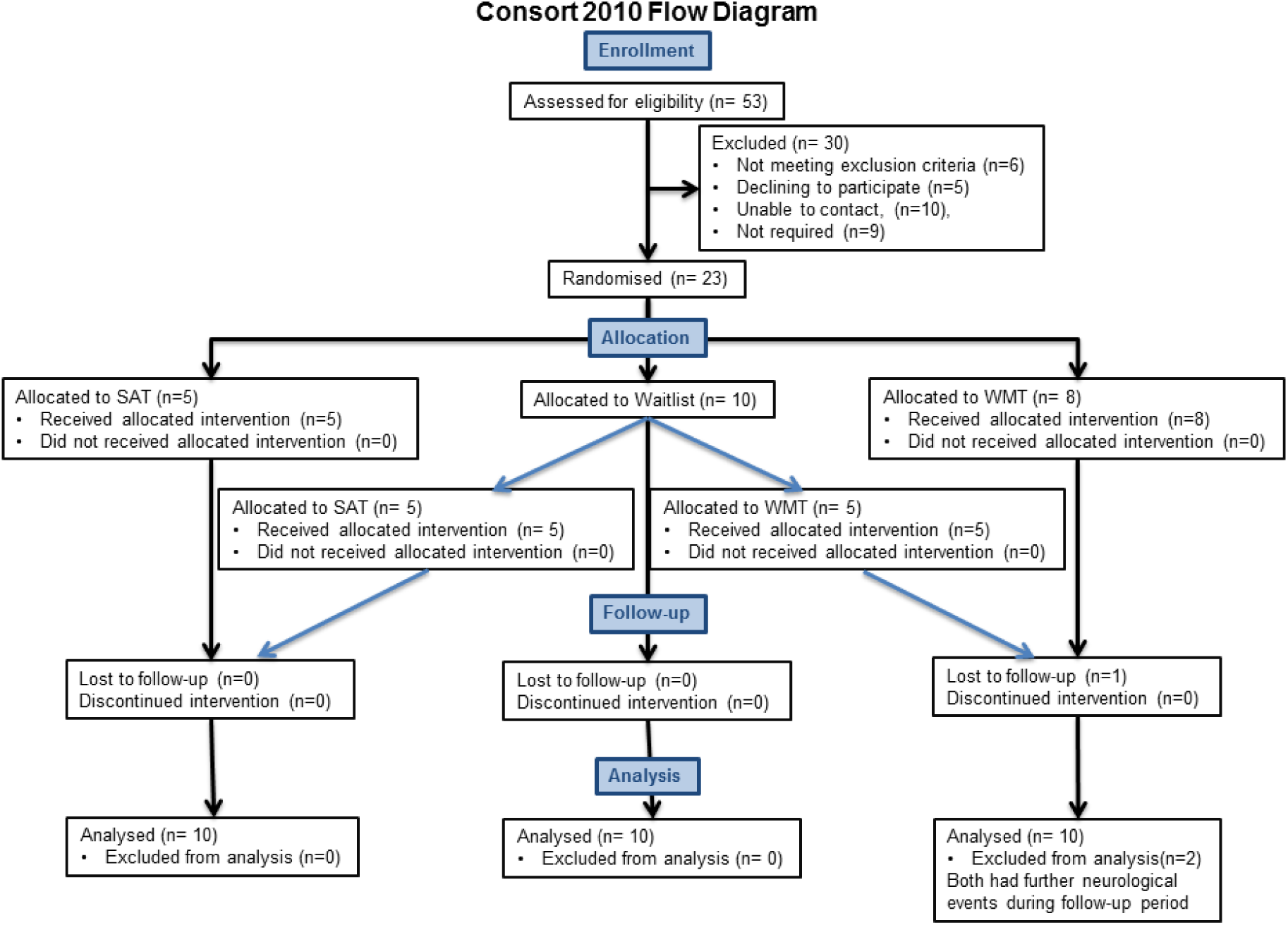
Consort flow diagram showing the progression of participant’s through the study

## Materials and methods

### Participants

Twenty-three participants from the Cambridge Cognitive Neuroscience Research Panel (CCNRP) gave informed, written consent for their participation in the study. The CCNRP is a database of volunteers who have historically suffered a brain lesion from various causes, and who have expressed an interest in participating in research. Twenty had right-hemisphere stroke, 2 left hemisphere stroke, and 1 had bilateral damage. All were chronic patients (mean time since injury 8.5 years, SD 4.7 years, range 7 months-17 years), aged under seventy-five years (mean 59 years, SD 10.6 years, range 28-74 years) and had no history of other neurological conditions. The recruitment of these patients with chronic lesions enabled us to collect outcome data on a wide-range of demanding, theoretically motivated, outcome measures to allow us to effectively assess the impact of training, minimising the issues of fatigue and daily fluctuations in ability often observed in acute patients. Patients were selected without knowledge of their behavioural difficulties, but on the basis that they had large lesions. Most had suffered middle cerebral artery (MCA) strokes, or had a lesion in areas of frontal and parietal cortices that have been linked to poor attention functioning (see Figure 2 for lesion overlays for the 10 patients for whom MRI scans were available). All had normal or corrected to normal visual acuity and sufficient language to comprehend and respond appropriately to the task demands and to provide informed consent. Although a number of the patients had substantial motor impairments they were able to make required responses (even if sometimes with their non-dominant hand) where these were required. The study was approved by the Cambridge Psychology Research Ethics Committee. Participants received a small honorarium for their time. The first patient entered the study on 15^th^ March 2013 and the last patient completed the study on 17^th^ September 2014.

**Figure 2.**
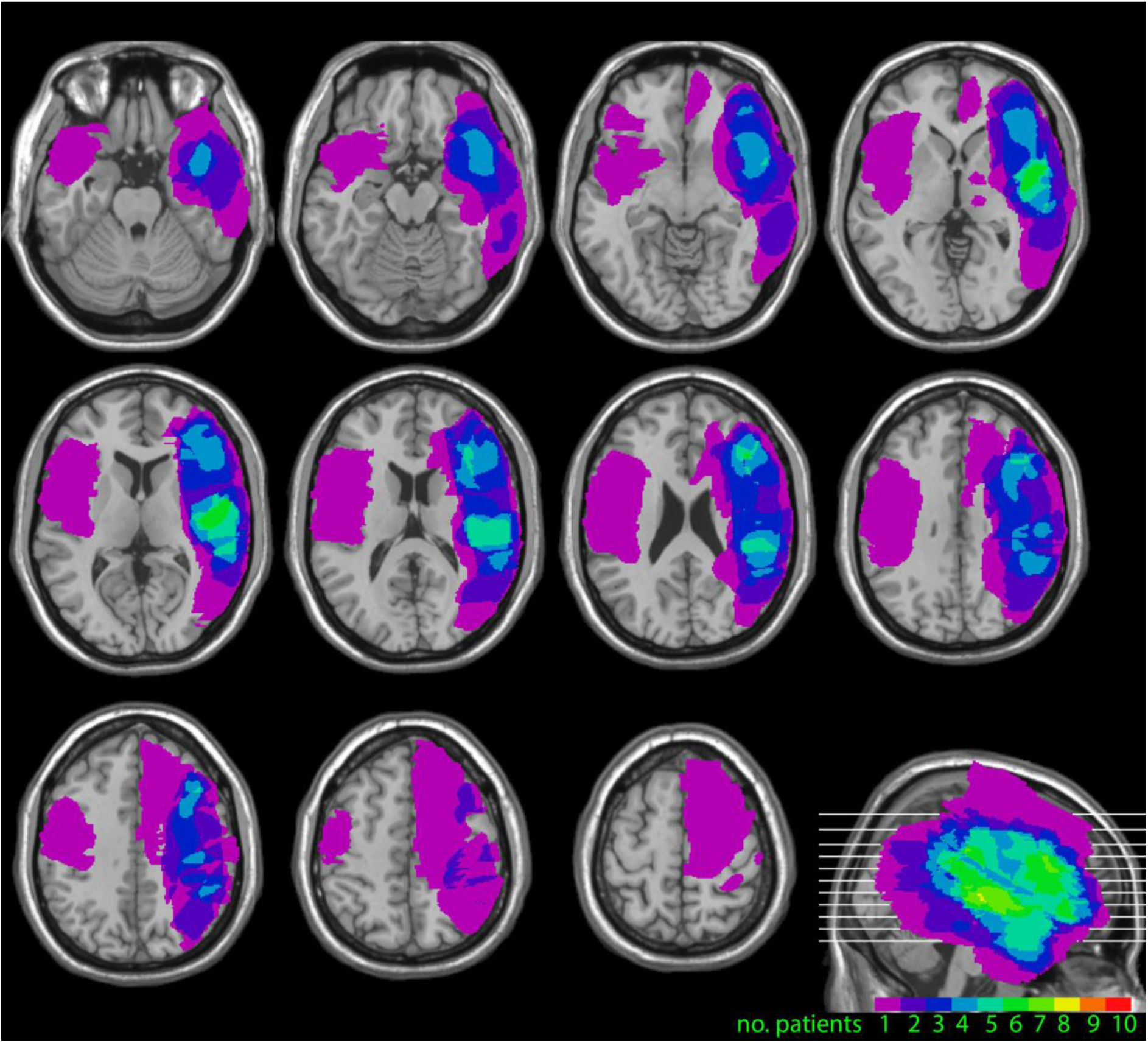
Lesion overlays for 10 of the 23 patients in the study, for whom scans were available. These show the foci of the lesions in frontal, parietal and temporal cortices predominantly in the right hemisphere.

### Procedure

Prior to their first assessment session participants were randomised into one of three conditions: WMT, SAT, or WL. At this time, the WL patients were further randomised to WMT or SAT to be completed at the end of the WL period. The entire randomisation sequence was completed by PP prior to the recruitment of the first participant and then subject numbers (and their corresponding conditions) were allocated by PP in a sequential order to participants as they became available without any prior knowledge of the individual.

After completing their initial assessment in their own homes, participants in training conditions were shown how to log in to the relevant websites and navigate through the tasks. They were asked to try and complete the training each weekday for the next 4 weeks (20 sessions). Participants were encouraged to get in touch with the project team if they experienced any difficulties accessing the tasks. In addition a weekly phone call was scheduled with a member of the research team in which participants could discuss any difficulties and their general progress. Because the research team received a log of each participant’s use of the programs, where repeated sessions were missed this could be brought up in the conversation, enquiring whether there were particular barriers, whether the participant had forgotten about the training and so on. If necessary we allowed longer than 4 weeks for participants to complete 20 sessions. All training sessions were completed in participants’ homes, for all but one on the participant’s home computer. WL patients also had a weekly phone call from the research team during their wait period in which they were asked similar questions about progress (but not about training). After the 2^nd^ assessment, when the Waitlist participants began training they received the same level of support described above.

### Pre-training assessment session

Participants received an extensive assessment of their cognitive profile and everyday functioning at each of the assessments. The measures focussed predominantly on attention and memory functioning and included both measures taken from basic science as well as those typically used to assess function in the clinic.

#### Background assessments

Participants completed a number of standard assessments including the Sloan Letter Near Vision Card (Good-lite Co, IL) to assess visual acuity, and the Tests for Colour-Blindness [31]. All patients had normal, or corrected to normal, visual acuity and all but one was found to have normal colour vision. The National Adult Reading Test (NART) [32] was used to estimate premorbid IQ and the Cattell Culture Fair Test [33] was used to estimate current fluid IQ.

#### Attention measures

*Partial and Whole Report TVA paradigm.* This test, based on tasks extensively used in many studies [14,15] and see [34] for review, was used to assess the attention parameters of spatial bias, attentional selection (α) and visual short-term memory capacity (VSTM) operationalised in TVA [21]. This task required participants to verbally report the identities of as many letters of a pre-specified target colour (either black or white) as they could from arrays of briefly presented targets and non-targets, whilst maintaining central fixation. Each trial followed essentially the same pattern. An initial red fixation cross flashed on and off a grey background at a rate of between 150 and 230 ms four times. An array of letters was then presented along with the fixation cross for 150ms before being replaced by the fixation cross alone until the experimenter had recorded all the participant’s responses and initiated the next trial. The arrays comprised of letters approximately 2 degrees by 3 degrees arranged in a circle approximately 10^º^ radius about the central fixation cross. Letters were selected at random from the set B,C,D,F,G,H,J,K,L,N,P,Q,R,S,T,U,V,X,Y,Z, and were presented in either black or white. Three basic types of array were presented; 1.*3 targets (3T)* Unilateral presentation of three letters (in the target colour) to either the left or the right of fixation. 2. *3 targets 3 non-targets (3T3NT)* Presentation of three target letters on one side of the screen with three non-targets (in the opposite colour) appearing on the other side of the fixation cross. 3. *6 targets (6T)* Presentation of six letters in the target colour, three to the left, and three to the right, of fixation. From these conditions, 3 separable attentional parameters (closely related to those defined in TVA, but using simplified formulae) were defined:

- *Absolute spatial bias*; the relative extent to which performance is preserved on a particular side of space in the presence of competing target information on the other side of space. To examine this, we compared relative reduction in performance between the *3T* and *6T* conditions for items presented on the left versus right sides of space, using the following formula:

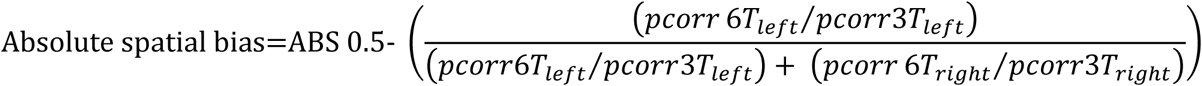

Where *pcorr* is the proportion of targets correctly identified in that condition.
- *Top Down Control* (α’); the extent to which distracting (non-target) information can be ignored. Here we examine where the performance in the *3T3NT* condition lies between the 3T condition and the 6T condition using the formula below. If participants have very good selection (lower values of α’), the non-targets should have relatively little impact whilst higher values of α’ indicate poorer attentional control.

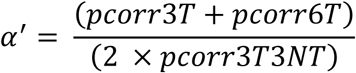
- *Visual Short-Term Memory Capacity* (K’); the maximum number of letters that can be reported from a brief display of letters. Following standard practice, we use probability mixtures of the maximum and 1- maximum performance. In this case *m* is the maximum number of letters ever reported (in the 6 target condition), and 6T_*m*_ is the number of trials in which the participant correctly reported *m* letters.

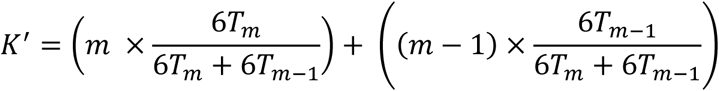
- *Variability.* In addition to the three traditionally measured TVA parameters an additional measure of participants’ variability in performance was derived. Variability in performance is thought to be indicative of poor sustained attention, which has also been linked to poor spatial awareness [35,36]. This was defined as the coefficient of variation (standard deviation divided by the mean) of correct letter reports from the *6T* condition.

Participants completed 4 blocks of 60 trials, two towards the start of the experimental session and two towards the end of the experimental session.

In addition to the TVA paradigm, participants completed 5 other computerised versions of attention measures that have either been used clinically or which have been shown to be sensitive to spatial bias in experimental studies. These were:

*Star Cancellation Task* [37]; a version of this well-known measure in which participants were asked to mark, using a stylus, all the small stars on a busy array of small and large stars and letters scattered across the screen as quickly as they could. Patients with spatial neglect have a tendency to miss a disproportionate number of targets from one side of the display.

*Line Bisection Test*; in which participants were asked estimate and mark the mid-point of seven lines, between 11.5 and 15.2 cm in length, presented either centrally or to the left or right of the screen. The bisections of patients with spatial neglect can deviate markedly from objective centre suggesting that their awareness of one end of the line is impaired [38].

*A Temporal Order Judgment Task*; in which two boxes appeared to the left or right of fixation either simultaneously or with a variable delay in their onset. The participants’ task is to judge which of the boxes appeared first. This version comprised 6 trials with simultaneous onsets and 2 trials with at each of 51ms,102ms, and 500ms onset asynchronies respectively. It has been reported that patients with left spatial neglect performing a similar task required the left target to appear up to 500 ms ahead of the right target before accurately reporting the order [39].

*A Lateral Reaction Time Task*; in which participants pressed a central button as soon as they detected a target that could appear either to the left or right of fixation. Fourteen targets were presented with variable inter-target intervals, equal numbers appearing on the left and right. Absent, disproportionately slow and variable response times to targets in neglected space have been reported [40].

*Slow and Variable Tone Counting Test*; in this variant of a test of sustained attention participants must attend to and count a variable series of tones separated by long and unpredictable intervals. Performance on this test, which has no spatial requirement, has been reported to be particularly poor in patients with persistent spatial neglect [41].

#### Working Memory measures

*Automated Working Memory Assessment (AWMA)* [42].

- *AWMA Dot Matrix Test*. In this computerised test a 4 × 4 grid was presented on the screen. The participant watched as a dot appeared at various locations on the grid and then recreated the sequence by pointing to the locations in the correct order. The test began with 2-location sequences and increased in sequence length until accuracy dropped below 50%.
- *AWMA Spatial Span Test*. Two abstract shapes were presented side by side on the screen. These could be identical or mirror images of one another, with the rightmost shape being presented in the upright position or rotated 120 or 240 degrees about the centre. With each presentation the participant had to determine whether the 2 shapes were the same or mirror images of one another. The shape on the right was always presented with a dot at one of three locations. At the end of a series of shape pairs the participant was asked to recall in order the locations of the dot on each pair. The test began with a single pair and increased the number of pairs until accuracy dropped below 50%.

#### Self-reported Everyday Function measure

*European Brain Injury Questionnaire (EBIQ)*[43]. This 63 item self-report questionnaire asks participants to rate their own function/symptoms over the preceding month. The items are grouped into nine broad categories; somatic symptoms, cognitive symptoms, motivation, impulsivity, depression, isolation, physical symptoms, communication issues and core symptoms, the latter being a global measure of disability.

### Training

The training batteries were internet based and completed in participants’ own homes. Following an initial induction they were completed without assistance from the research team. The batteries shared some essential core features, namely: that they were adaptive and therefore designed to keep patients working at their maximal ability, and that trial by trial feedback was given for both learning and motivational purposes.

#### Working Memory Training (WMT)

The adaptive version of the commercial Cogmed^**™**^ Working Memory Training (Pearson; for full details see www.cogmed.com/rm) was used.

Participants attempted 15 trials of 8 tasks in each session, covering both verbal and visuo-spatial working memory. Following the standard set-up, three of the twelve tasks in the battery were presented in every session, with the rest of each session being made up of five of the remaining nine tasks. Most participants completed a session of training in approximately 30-50mins.

#### Selective Attention Training (SAT)

This training was designed by the research team and programmed in Flash using Adobe Flex Builder 3. They were deployed via a custom website (https://www2.cbstrials.com) developed in Ruby on Rails. The training consisted of five time-limited tasks designed to improve selective attention, comprising:

#### Aliens Task

In each trial an onscreen array of cartoon aliens appeared, one of which was designated as the target. The participant’s task was to decide as quickly as possible whether another of the aliens was an exact match to the target, indicating a match/mismatch response by mouse clicking onscreen buttons (S1 Figure). All aliens were comprised of a combination of a head part, a body part and legs selected from four prototype heads, bodies and legs. These could vary along parameters such as the texture and thickness of arms and legs, number of eyes, the presence/absence of tail, hairstyles and clothing. With correct responses, task difficulty was increased by increasing the number of aliens in the array and their similarity to the target (requiring increasing attention to small distinguishing details). As with all of the SAT tasks, auditory and visual feedback was given for correct (a large green tick and a bell) and incorrect (a large red cross and a buzz) responses, progress was indicated by an onscreen thermometer and the remaining time for the task indicated with an onscreen digital clock. The duration of Aliens in each training session was 3 minutes.

#### Visual Search

In each trial an abstract shape was presented on the screen for a few seconds. It was then replaced by an array of objects (S2 Figure) and the participant was asked to judge whether any exactly matched the original shape. Difficulty was manipulated by increasing the similarity of the objects to the target along dimensions of shape, size, colour and texture. The task was played for 4 minutes on each training session.

#### Jigsaw

At the top of the screen two or more red boxes were shown each containing a distinct pattern or object (e.g. one with blue and white stripes, the other with an inverted grey triangle). In the lower part of the screen four or more white boxes also appeared, each with patterns or shapes. The participants’ task was to decide whether the elements of each red box were present in the white boxes such that the ‘jigsaw’ could be made from these pieces (S3 Figure). Difficulty was manipulated via the similarity of the elements in the boxes to the target configurations, the number of red boxes that needed to be matched and whether the elements in the white boxes needed to be mentally rotated to make up the patterns. The task was played for 4 minutes on each training session.

#### Rotations

This test required participants to take in the spatial relations between a series of shapes and then mentally rotate this image to judge whether it would match a second image (S4 Figure). In each trial two large squares were presented, each containing one or more smaller green or red squares (in effect, filled cells of an invisible identical grid). The participant’s task was to indicate whether rotation of one large square and its elements would make it identical to the other. Difficulty was manipulated by the number of elements within the squares, the degree of rotation required and the similarity of the two squares (e.g. the elements being in very different locations compared to only one of many elements differing). Participants played the task for 3 minutes in each training session.

#### Button Sorting

On each trial of this set-shifting task a shape was presented upon which the participant was asked to make a speeded judgment based on a rule also presented on the screen (S5 Figure). If the rule was ‘shape’ the participant had to indicate whether the shape most closely resembled a circle or square by clicking on an arrow pointing to one of two reference shapes (circle and square) that were coloured red and yellow (the color was irrelevant to the ‘shape’ rule). If the rule changed to ‘color’ the participant had now to click on the arrow pointing to the correct color and ignore the shape of the reference. Difficulty was increased by morphing the shapes in the direction of the alternate category (e.g. increasingly rounding off the square) and making the colors increasingly similar. Participants played the task for 4 minutes on each training session.

### Post-training assessment

These sessions comprised the same tasks as the pre-test session without the background assessments. Sessions were scheduled to occur within 2 weeks of completing the online training, or the case of the WL, 4-6 weeks from their initial assessment.

The study was not blinded, and as an exploratory trial based on samples sizes reported in previous training studies on stroke patients, showing positive effects [23].

The study was not prospectively registered as a clinical trial as it was set up as an exploratory study to investigate whether training may be beneficial in a group of individuals who might be expected to have reduced attention, to see whether such an approach may have potential in a clinical sample. The primary interest of the study was to look at changes on experimental measures of attention and working memory. Of secondary interest was whether any improvements would transfer to changes in everyday life. All data collection was completed prior to the NIH publishing its definition of a clinical trial on 23^rd^ October 2014. The authors confirm that all ongoing and related trials for these interventions are prospectively registered.

## Results

Complete data were analysed from twenty of the twenty-three participants. Of those whose were omitted, two were removed having suffered subsequent neurological events between initial assessment and final assessment and one had to drop out owing to family circumstances. One-way ANOVAs were carried out to see whether the 3 groups differed on background measures. No significant differences between the groups were observed for age (F(2,29)=0.57), time since injury (F(2,29)= 1.61), visual acuity (F(2,29)=1.20), NART (F (2,29)= 1.37) or Cattell (F (2,29)= 1.37) suggesting the groups were well matched.

### Training

#### 1 Compliance

Compliance with the training program was generally good. All patients who started the training completed the study and on average the WMT group completed 19.8 of the intended 20 sessions (range 18-20 sessions) whilst the SAT group completed 20.2 sessions (range 18- 23). Patients were in regular contact with the research team (by phone or occasionally email) over the course of the training period. Participant feedback regarding the training was generally very positive. Nine participants requested to continue with training following their final assessment.

#### 2 Improvement on training tasks

Mean performance by session data (collapsed across the three continuous Cogmed tasks, or all SAT tasks) for each of the training batteries are shown in Figure 3. Polynomial equations (y= x^2^ + x+ c) were fitted for each participant separately. These provided better fits than logarithmic fits and allow us to determine parameters including δy (the improvement in performance), δx (the number of sessions to maximal performance) and δy/δx (the average rate of improvement to asymptote). Polynomial fits were good, with a mean r^2^ of 0.76 (range 0.56-0.87) for the WMT group and 0.91 (range 0.68- 0.99) in the SAT group, for all but three patients. Data from these three patients with were therefore excluded from analyses of training gains, but not from analyses of outcome measures. Generally, the two training conditions appear to show a similar improvement profile with maximal performance achieved after 15.6 sessions (range 11.8- 18.8) and 16.6 sessions (range 13.25-23.54) for the WMT group and the SAT group respectively. Direct comparison of the improvements in the two training conditions were precluded by the different scales used. Nonetheless, the WMT group showed an average improvement of 2.7 items (range 1.2- 4.6) whilst the SAT group improved by 11.7 points (range 7.9- 16.1) indicating that all participants were able to demonstrate improved performance with training.

**Figure 3.**
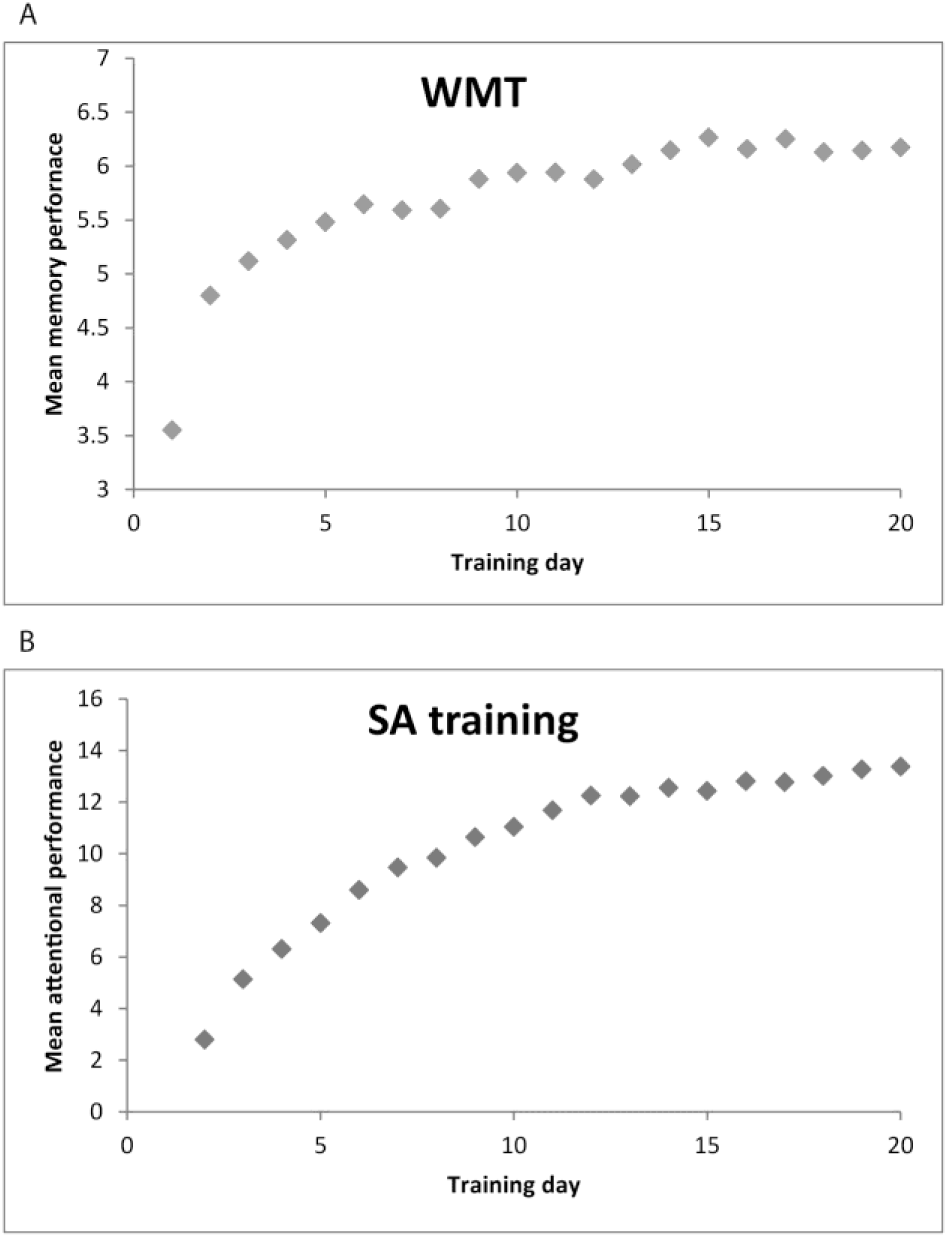
Average performance on the training tasks over the twenty days of training for (A) WMT and (B) SAT groups respectively

#### 3 Training transfer

Having demonstrated that participants generally improved on the training tasks, we next established whether these improvements transferred to other cognitive tasks and measures of disability. For all subsequent analyses a regression approach was used to examine whether post-test performance was influenced by training group (WL, WMT or SAT) whilst adjusting for pre-test performance. Coding dummy variables allowed us to compare the effects of the interventions (i.e., WMT compared to WL and SAT, and SAT compared to WL and WMT) in a single analysis. This regression approach is a stricter test of training gains than standard ANOVAs because interactions in the ANOVA can be at least partly driven by pre-training differences. For completeness repeated measures ANOVAs were also carried out, these showed essentially the same pattern of results as the regression analyses and are not reported here. In addition to the regression approach, paired sample t-tests were carried out to examine whether post-scores differed significantly from pre-scores for each of the groups. For several transfer measures, Figure 3 shows post-test score minus pre-test score for each of the three groups.

#### Measures of attentional functions

Turning first to spatial bias (see Figure 4a), the regression indicated that between them, ‘pre-test score’ and ‘experimental group’ predictors explained 75.5% of the variance (R^2^ =0.76, F(3, 26)= 26.74, p<0.001). Whilst as might be expected, ‘pre-test score’ was a significant predictor (β=0.89, p<0.001), SAT (compared to a combination of WL and WMT) was also a significant predictor (β=-0.41, p=0.002), whereas no such effect was seen for WMT (compared to a combination of WL and SAT) (β=-0.18, p=0.15). In addition to this, paired samples t-tests indicated a significant change in bias score between pre and post testing in the SAT group (t=-3.03, df=9, p<0.05), no such change was observed in either the WMT group (t=-1.27, df=9, p=0.24), or WL (t=1.44, df=9, p=0.19). Thus SAT alone appeared to have a beneficial impact on spatial awareness.

**Figure 4.**
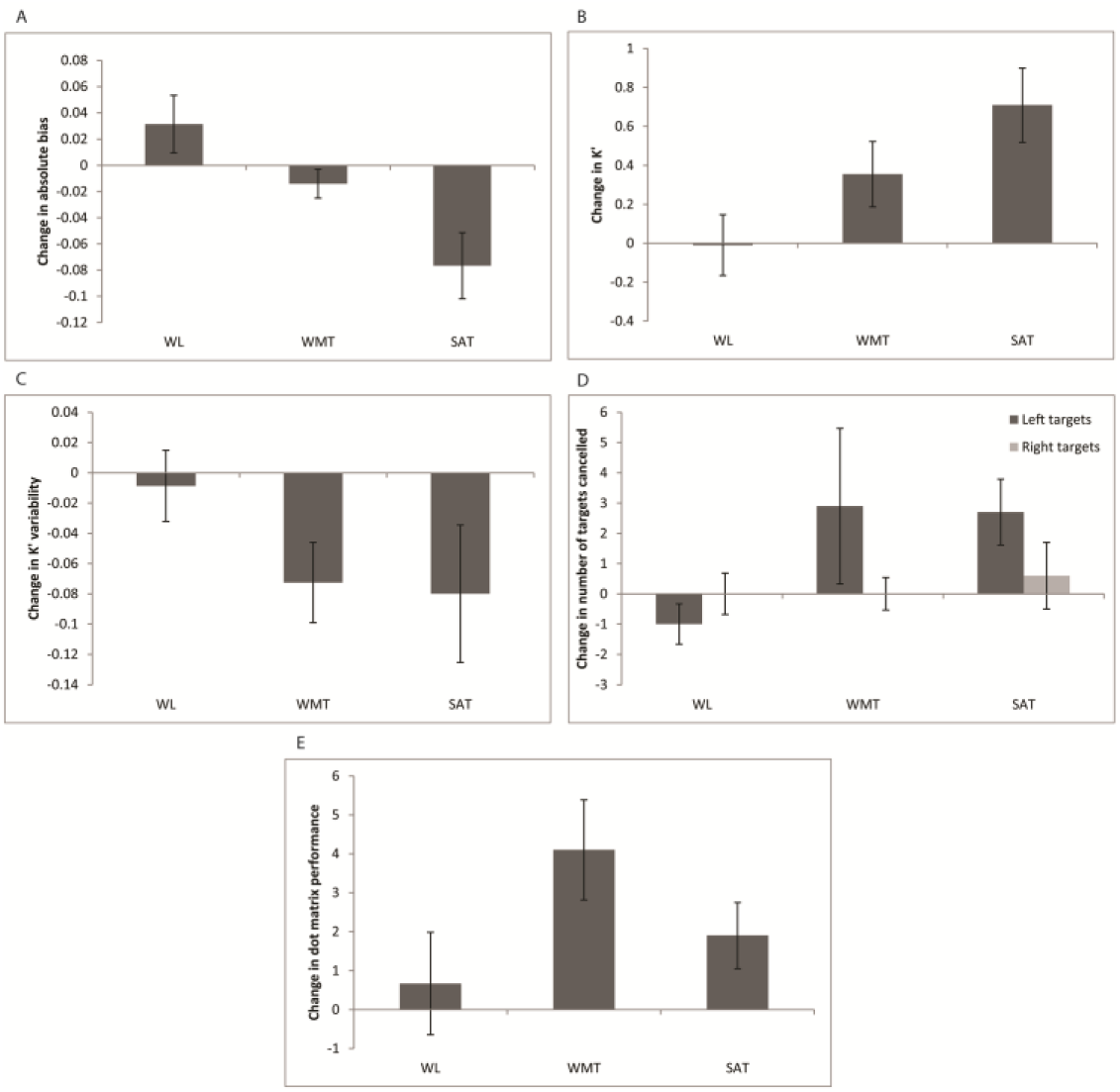
Mean (± S.E) change in performance from pre-test to post-test for the experimental measures in each of the groups. Plots show performance for; A. change in TVA absolute bias, B. change in K’, C., K variability, D. change in number of targets cancelled by side on the star cancellation task and E. change in Dot Matrix performance.

As our SAT was focussed on improving selective attention it may be expected that any attentional affects on awareness may stem from improvements in top-down control (α’). However, the regression indicated that between them, ‘pre-test score’ and ‘experimental group’ predictors explained less variance than we saw with spatial bias (R^2^=0.57, F(3, 26)= 11.54, p<0.001) and ‘pre-test score’ was the only significant predictor (β=0.74, p<0.001).

The K’ capacity measure might have been expected to have been influenced by both WMT and SAT training. The regression indicated that between them, ‘pre-test score’ and ‘experimental group’ predictors explained 68.7% of the variance (R^2^ =0.69, F(3, 26)= 19.02, p<0.001) (Figure 4b). In line with our prediction it was found that ‘pre-test score’ (β=0.77, p<0.001), SAT (β=-0.38, p<0.01), and to a lesser extent WMT (β=-0.28, p<0.05) were all significant predictors of post-test score. Paired samples t-tests indicated that K’ values were significantly improved in the SAT (t=3.73, df=9, p<0.01) group post training, an effect that reached near significance in the WMT (t=2.11, df=9, p=0.06) group, but was absent in the WL(t=0.06, df=9, p=0.95).

It is worth noting that K’ and spatial bias might not be independent. To take an extreme example, if a participant reports all 6 letters, spatial bias must be zero. To address this potential non-independence we re-ran the spatial bias regression including ‘pre-post-test K’ change’ as a predictor of pre-post-training bias change. This indicated that, between them, ‘pre-test score’, ‘change in K’ and ‘experimental group’ predictors explained 76.1% of the variance (R^2^ =.761, F(25,29)= 19.89, p<0.001). Whilst ‘pre-test bias’ (β=0.87, p<0.001) was a significant predictor, critically, ‘change in K’ was not (β=0.09, p=0.49). Importantly, despite this very stringent test, SAT (β=-.36, p<0.05) remained a significant predictor, but not WMT (β=-.16, p=0.20). This strongly suggests that the effects of improved spatial bias following SAT were not simply an artefact of improved capacity.

We were also able to use 6T variability (Variability) as a measure of the consistency with which attention was maintained (Figure 4c). Despite our training not being specifically designed to develop this skill, ‘pre-test score’ and ‘experimental group’ predictors still explained 36.8% of the variance (R^2^ =0.37, F(3, 26)= 5.05, p<0.01) in post-test Variability. This is driven by both ‘pre-test score’ (β=0.51, p<0.005), and WMT (β=-0.37, p<0.05). Here, no effect of SAT was seen (β=-0.20, p=0.28). Paired samples t-tests indicated that Variability was significantly reduced in the WMT (t=2.73, df=9, p<0.05) group post training, but such reduction was not observed in the SAT (t=1.76, df=9, p=0.11) or the WL(t=0.37, df=9, p=0.72) groups.

Analysis of the standard clinical measures of attention, star cancellation, line bisection, prior entry, lateral reaction time and line bisection were carried out despite most patients showing no significant clinical impairments on these tasks at pre-test (patients had chronic lesions and were selected on the basis of lesion location rather than clinical symptoms). As exemplified by the star cancellation data (see Figure 4d) an encouraging pattern of results was observed post-training, with increased awareness on left sided items, but this failed to reach statistical significance.

#### Working Memory Measures

Change in performance on a measure of visuo-spatial capacity, the AWMA Dot Matrix task is shown in Figure 4e. The regression indicated that between them, ‘pre-test score’ and ‘experimental group’ predictors explained 62.8% of the variance (R^2^ =0.63, F(3, 26)= 14.09, p<0.001). In line with our prediction ‘pre-test score’ (β=0.73, p<0.001) and WMT (β=0.30, p<0.05) were significant predictors of post-test score, while SAT was not (β=-0.09, p=0.59). Paired samples t-tests indicated that the number of locations correctly recalled was significantly increased in the WMT group (t=3.19, df=9, p<0.05) post-training, an effect that reached near significance in the SAT group (t=2.24, df=9, p=0.05), but was absent in WL (t=0.51, df=8, p=0.63). Performance on the Spatial Recall task of the AWMA did not vary by training condition in the same way. Although a significant regression (R^2^ =0.55, F(3, 26)= 9.03, p<0.001) was observed, the only significant predictor of post-test performance was ‘pre-test score’ (β=0.70, p<0.001) with neither WMT (β=0.24, p=0.15) nor SAT (β=0.04, p=0.85) acting as significant predictors. Despite this, paired-sample t-tests indicated that the WMT group showed a significant improvement in performance between pre- and post-test (t=2.49, df=9, p<0.05) whereas neither SAT (t=0.48, df=8, p=0.65) nor WL (t=0.15, df=8, p=0.88) showed such effects.

#### Measures of disability

Changes in disability rating for each of the domains of the EBIQ are shown in figure 5. To limit the number of statistical tests conducted, formal analysis was limited to the two most pertinent domains; core symptoms (a global measure of impairment) and cognitive symptoms. Turning first to core symptoms, regression indicated that between them, ‘pre-test score’ and ‘experimental group’ predictors explained 66.4% of the variance (R^2^ =0.66, F(3, 26)= 19.02, p<0.001). In line with our prediction, ‘pre-test score’ (β=0.65, p<0.001), WMT (β=-0.61, p<0.001), and SAT (β=-0.58, p<0.001) were all significant predictors of post-test score. Paired samples t-tests indicated that core symptoms were significantly reduced post-training in both the WMT (t=-5.42, df=9, p<0.001) and the SAT (t=-4.38, df=9, p<0.005) groups, but not in the WL(t=0.77, df=9, p=0.46). In a similar manner, a regression analysis indicated that ‘pre-test score’ and ‘experimental group’ predictors explained 55.7% of the variance (R^2^ =0.56, F(3, 26)= 10.92, p<0.001) in post-test cognitive symptoms. Paired samples t-tests indicated that cognitive symptoms were significantly reduced post training in both the WMT (t=-5.78, df=9, p<0.001) and the SAT (t=3.41, df=9, p<0.005) groups, but not in the WL (t=-1.15, df=9, p=0.28) group.

**Figure 5.**
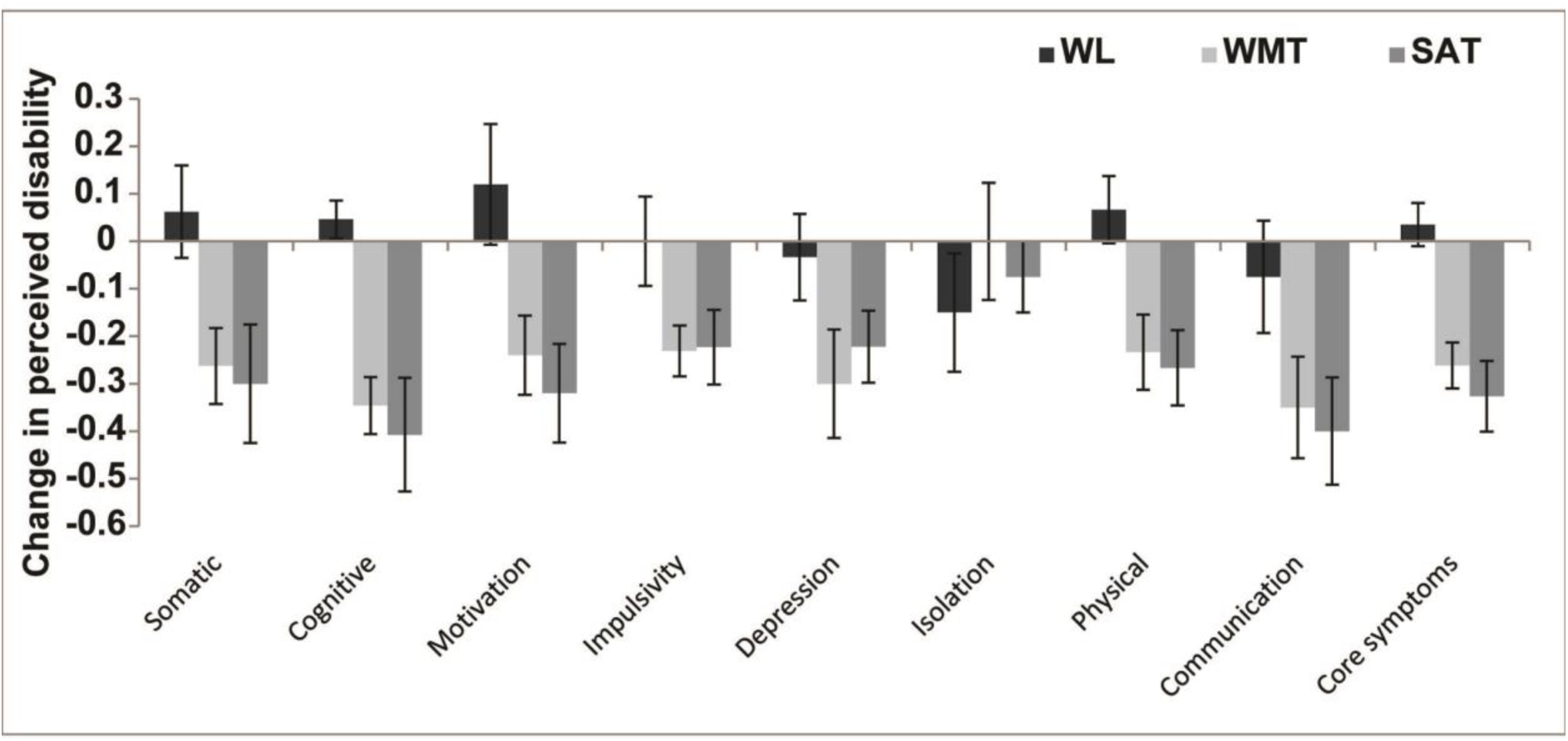
Change in self-reported disability as measured by the EBIQ from pre-test to post-test.

#### What predicts reductions in disability?

A key question is whether self-reported improvements were related to objective changes in cognitive function. A regression using ‘pre-test core symptoms’, ‘change in absolute bias’, ‘change in K’, ‘change in Dot Matrix performance’ and ‘change in Variability’ as predictors of pre-post test change in core symptoms indicated that these variables explained 63% of the variance (R^2^ =0.63, F(5, 28)= 7.74, p<0.001). As may have been expected, ‘pre-test core symptom score’ was a significant predictor (β=0.44, p=0.006) of change in reported core symptoms. In addition both ‘change in absolute bias’ (β=0.37, p=0.016) and ‘change in Variability’ (β=0.33, p=0.03) significantly predicted change in core symptoms. This was not true for changes in K’ (β=0.03, p=0.83) or Dot Matrix performance (β=-0.20, p=0.15).

#### How does training improvement influence transfer?

The specificity of some of the improvements in working memory and attention tasks may be indicative of task or domain specific training. If this were the case, we might expect that the extent of the training gain would be predictive of the extent of improvement in closely related outcome tests and less predictive of change on more divergent measures. To compare training rates in the two training groups we standardised the rate parameter δy/δx, (Zδy/δx), and then generated in interaction term based on the product of the de-meaned group and the newly standardised rate parameter. Regressions were carried to examine significant predictors of change in performance between pre- and post-test on the basis of: ‘pre-test score’, ‘intervention’ (WMT vs. SAT), ‘rate of improvement on training’ (Zδy/δx), or the ‘interaction between group and improvement rate’ (Gp*Zδy/δx). Turning first to change in Dot Matrix performance, the regression indicated that between them, ‘pre-test Dot Matrix score’, ‘intervention’, Zδy/δx and Gp*Zδy/δx predictors explained 70.6% of the variance (R^2^ =0.71, F(4, 12)= 7.21, p<0.005). ‘Training type’ (β=0.35, p<0.05), ‘training improvement’ (Zδy/δx;β=-0.61, p<0.05), and ‘Training type x training improvement’ (Gp*Zδy/δx; β=0.58, p<0.005) were all significant predictors of change whilst ‘pre-test Dot Matrix performance’ was not.

As might be expected from this finding and shown in Figure 6a, the patients who showed the biggest WMT training also showed the biggest improvements on the untrained though similar Dot Matrix tasks, whereas the extent of SAT training gain did not influence Dot Matrix improvement. The pattern of results was markedly different for attentional measures. For both ‘change in absolute bias’ (see Figure 6b) and ‘change in K’’, the regressors failed to significantly predict variance in change scores. As Figure 6b demonstrates, within the SAT group there was virtually no difference in change in bias scores between those who made small and large training gains. Changes in Variability and EBIQ core symptoms were significantly predicted by ‘pre-test core symptoms’, ‘training type’, ‘training gains’ and ‘training gains x training type interaction’ (R^2^ =0.62, F(4, 12)= 4.91, p<0.05 for K variability, and R^2^ =0.70, F(4, 12)= 6.88, p<0.005 for the EBIQ core symptoms). However, in both cases ‘pre-test score’ was the only significant predictor of change (β=0.74, p<0.005 for K variability and β=0.89, p<0.001 for EBIQ core symptoms) and as Figure 6c shows, for both the WMT and SAT groups, similar reductions in core symptoms were reported by those with relatively small and large training gains.

**Figure 6.**
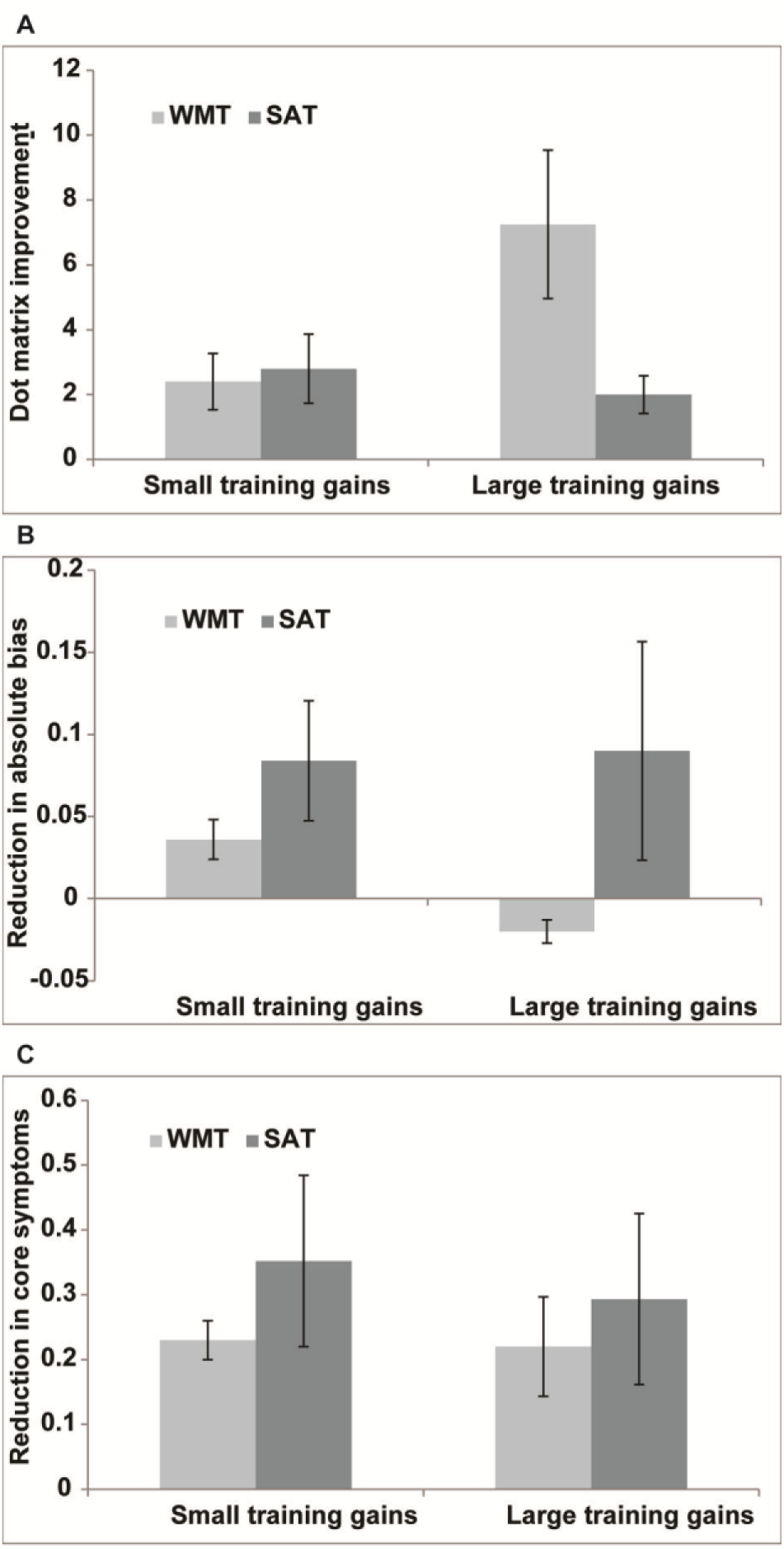
Mean (±S.E.) changes in performance for patients who made small and large training gains (based on a median split) as a function of training type. A. Dot Matrix task. B. Absolute spatial bias. C. EBIQ core symptoms.

## Discussion

This exploratory proof-of-concept study examined whether two forms of training aimed at improving attention would lead to improvements in untrained outcome measures and self-reported disability in individuals with chronic brain lesions. A good level of compliance and approximately equivalent exposure to training between the groups allowed us to realistically assess objective benefits and examine whether the distinct training programmes had distinct and/ or generic effects. Whilst we acknowledge the relatively modest sample sizes should be kept in mind, the study provides preliminary evidence which can inform the development of future studies in clinical samples, for whom, the rigours of the extensive experimental assessments may not be appropriate.

### Distinct training effects

Participants who trained on the commercially available WMT showed greater improvements than SAT and WL participants on the Dot Matrix working memory outcome measure. Such ‘near transfer’ of training gains has been observed in other populations [44,28]. Indeed improvement in Dot Matrix performance was strongly predicted by the extent of training gains made by the WMT group, pointing to learning of specific task related strategies for spatial span. Given that improvements beyond a narrow training context are a pre-requisite for any likely transfer to everyday activities, but that previous work [45] has shown that even these specific training activities do not always transfer to even closely related tasks, the current finding is encouraging. It suggests that patients can effectively apply memory related strategies developed in training to very closely related tasks. There was some evidence of improvement on the more complex Spatial Recall task in the WMT group; however, WMT was not found to disproportionately influence post-test Spatial Recall relative to SAT and WL suggesting that transfer effects to more distantly related memory tasks may be relatively small, and questioning the generic advantage of such interventions.

Perhaps of greater interest, however, are the improvements seen in the SAT participants across a wider variety of attentional parameters, on paradigms that appear quite different from the trained tasks and which are less dependent upon the extent of the improvements made on the training tasks. SAT participants disproportionately improved, relative to those in WMT and WL conditions, in their ability to take in more information ‘at a glance’ (K’) from brief displays and in the reduction of spatial bias (of which there were also hints in the clinical measures). The first is plausibly linked to differences in the training tasks. In WMT participants typically monitor sequences in which a single event occurs at any one time. In contrast, solving the problems presented in SAT involved taking in an increasing amount of visual information. Encouragingly, improving this capacity in a particular context during training led to attentional improvements that participants were able to effectively utilize in different contexts. It is possible that this practised distribution of attention also underpinned the reduction in spatial bias. However, at least in the context of unilateral spatial neglect, even explicit training of visual scanning *per se* has often proved of limited generalised efficacy [46]. Another possibility, alluded to in the introduction, is that the reduction in spatial bias is a consequence of generally improved attentional ‘tone’ – a relatively alert state in which relevant information from across space is better prioritized. It has previously been shown that fluctuations in alertness from stimulant medication, loud tones, time-on-task, and sleep onset can impact on patients’ and healthy participants’ relative awareness for information on the left and right sides of space [35,18,20]. Anecdotally, both SAT and WMT patients appeared more awake and engaged after 4 weeks of regular, monitored cognitive activity with direct feedback. Whilst improvements in our proxy of alertness (TVA performance variability) were actually greater on average for the SAT than WMT groups, the substantial variability across SAT participants meant that this change failed to reach statistical significance (Figure 4c). Variability scores and change-in-variability scores tend, by their nature, to be somewhat unreliable as noise from the underlying measures is summed and further work is required in operationalizing ‘alertness’ and understanding mechanisms of change.

### Generic effects

In addition to improvements that were specific to WMT or SAT, more general positive effects of training were observed, particularly a marked reduction in self-reported disability across both training groups. Importantly these reductions were significantly influenced by improved spatial bias and reduced variability in performance, suggesting a link of self-perception to measureable changes in attentional functions. Interestingly, improvements in WM span did not significantly influence self-reports in the same way. If this finding is replicated one possibility is that SAT practice indeed produces deeper or faster generalised changes for everyday cognition than WMT. Various accounts can be proposed for such an effect. Firstly, previous studies have suggested that poor attentional functioning is particularly associated with high levels of disability and poor outcome [7,8]. Hence change in these capacities may also produce more generalised effects. Secondly, gains in WMT may be disproportionately achieved via strategy development (see similar findings in Alzheimer’s Disease, [47]) rather than underlying capacity, and as our data on transfer to other WM tasks suggest, strategy may be less easily generalised to different contexts. Other possibilities are that the greater effects of SAT on everyday function relates to the intentional recruitment of predominantly right-hemisphere patients in our sample (for whom attention deficits may be the primary cause of issues with activities of everyday living), or SAT being perceived as more relevant and hence being more influential over self-report.

Somewhat unexpectedly, as discussed, WMT was linked with significant reductions in performance variability on the TVA attention measures. If reliable, such transfer to a seemingly unrelated task is particularly striking given some previous literature suggesting WMT gains are restricted to near transfer to very similar span tasks [28]. However, it is not implausible to imagine how repeated practice of monitoring increasing sequences of spatial of verbal material for subsequent recall, during which even a brief lapse could prove disastrous for the entire trial, could progressively shape such consistent engagement. Along these lines, studies in the Behavioural Activation literature (encouraging patients to schedule and participate in rewarding, stimulating activity) suggests that engagement in mentally stimulating activities may help to improve alertness [48]. It is possible, therefore, that providing a daily structure within which patients were helped to focus on a cognitively demanding task for a relatively prolonged time may be sufficient to help improve alertness, perhaps irrespective of the precise demands of that training.

### Appropriateness of home-based computer training for patients

The potential efficacy of cognitive training batteries to improve outcome has been vigorously debated in recent times, in both healthy adults and the developmental literature, with many suggesting that improvements may be short lived and fail to generalise to meaningful improvements in everyday functioning [49,29,30]. Our data showing reductions in self-reported disability related to improvements in attentional functions suggest this may not be the case in stroke patients. As discussed, there are plausible reasons why this population may benefit from training in a way that the developmental population may not. Providing some structure, focus and stimulation, as well as clear feedback to help them learn, may be critical to reductions in disability. Whilst these aspects of training are already in place in a school environment, many patients receive little input from clinical services and lack structure or focus to their day. Along these lines, positive effects of online training on both cognitive function and activities of daily living have been observed in healthy older adults [50].

The success of any intervention is dependent not only upon the potential for improvement following treatment, but also upon how practical and tolerable it is for patients. Here patients’ ability to cope with navigating to websites, logging in etc. was good and attitudes to both interventions were generally positive, with a good proportion of patients feeling it was worthwhile continuing after the study. In accordance with this, despite the time commitment of the study, drop-out rates were very low. A caveat is that this sample was recruited from a panel of individuals who have already indicated that they are motivated to take part in research. It remains to be seen whether such good compliance would be seen in an unselected population of stroke patients.

### Implications and future directions

The results so far indicate some specific effects of the two types of training and some generally positive effects from both compared to WL. The specific training effects are well controlled in terms of exposure to training, interaction with the experimenter and the knowledge of being engaged in training hypothesised to be helpful. However, interpretation of the more general effects, is limited by reduced stimulation in the WL and potential expectancy effects. To a degree this is offset by the finding that reductions in spatial bias and improved K’ variability over the course of the study predicted changes in self-reported disability, suggesting that improvements in attentional functioning could be key to reducing disability. Of course the reverse causality also remains a possibility. An active and plausible control condition hypothesised not to be beneficial is required to clarify these issues.

It is generally accepted that the majority of spontaneous recovery occurs within the first six months after stroke [51,52] and it is therefore perhaps surprising we saw such extensive training effects on average 8 years post-injury. Whether training gains in the chronic phase may be more attributable to strategy development than underlying recovery remains an important topic of investigation.

In summary, our study provides evidence that cognitive training is feasible in stroke patients, and can lead to both specific improvements in cognitive functions and more general reductions in self–reported disability. Further work is required to examine whether such effects can be replicated in a larger sample.

## Acknowledgements

This work could not have been carried out without the willingness and effort of our patients and their families.

## Supporting information

**S1 Figure.**
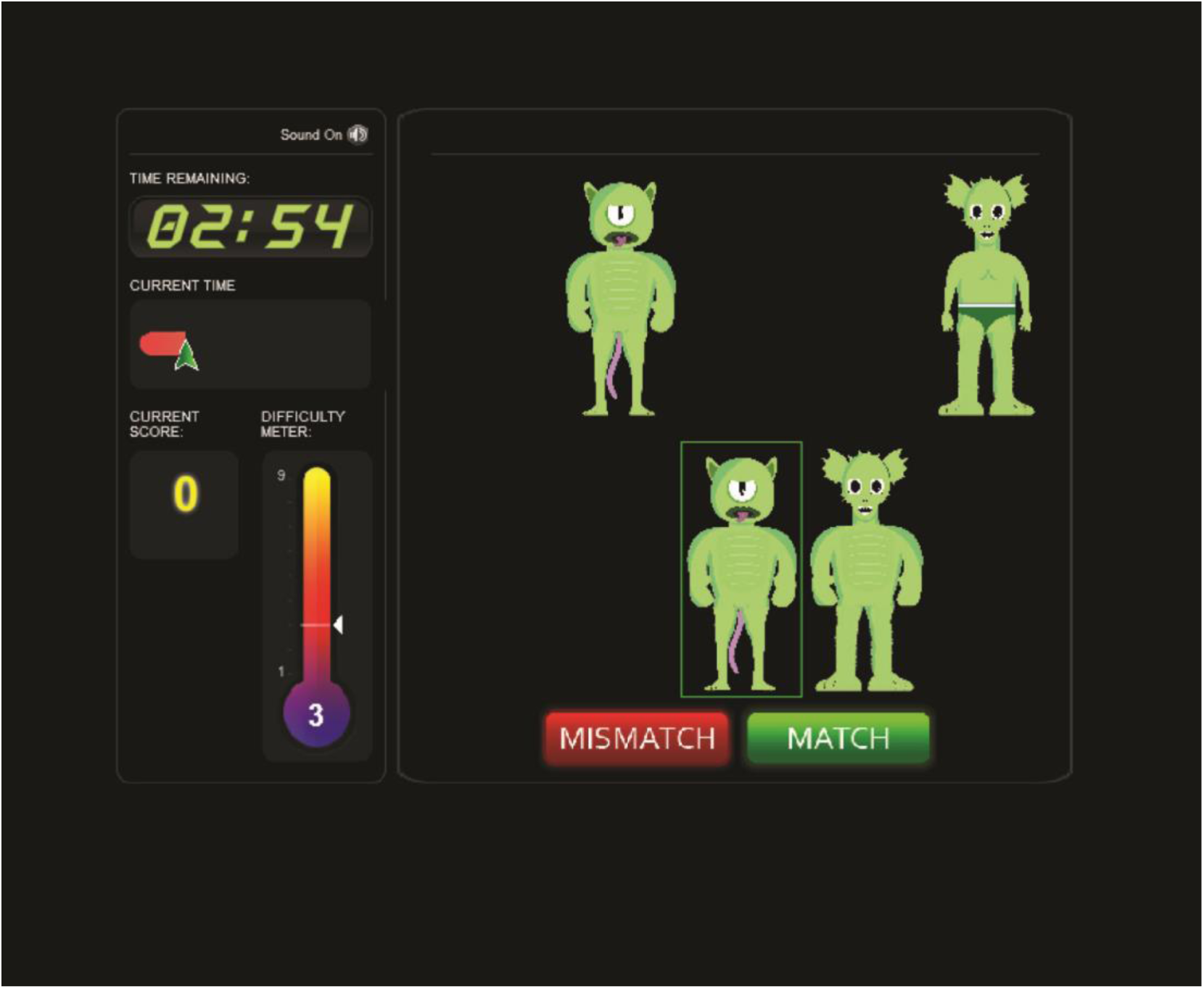
An example screen from the aliens task. This example shows a ‘match’ trial with the highlighted alien matching the on in the top left corner.

**S2 Figure.**
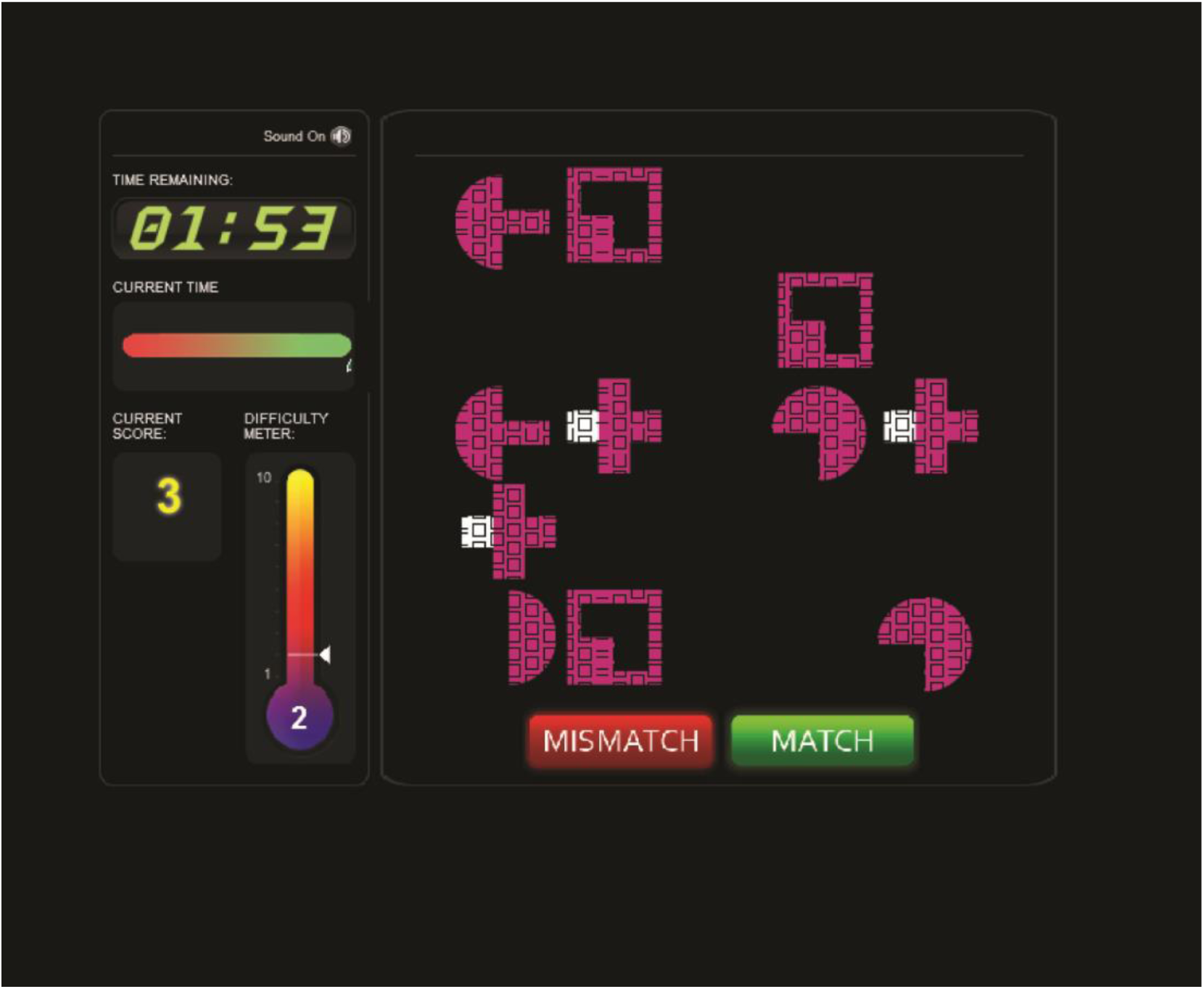
An example array from the visual search task, participants had to say whether an exemplar they had just seen was present in this array.

**S3 Figure.**
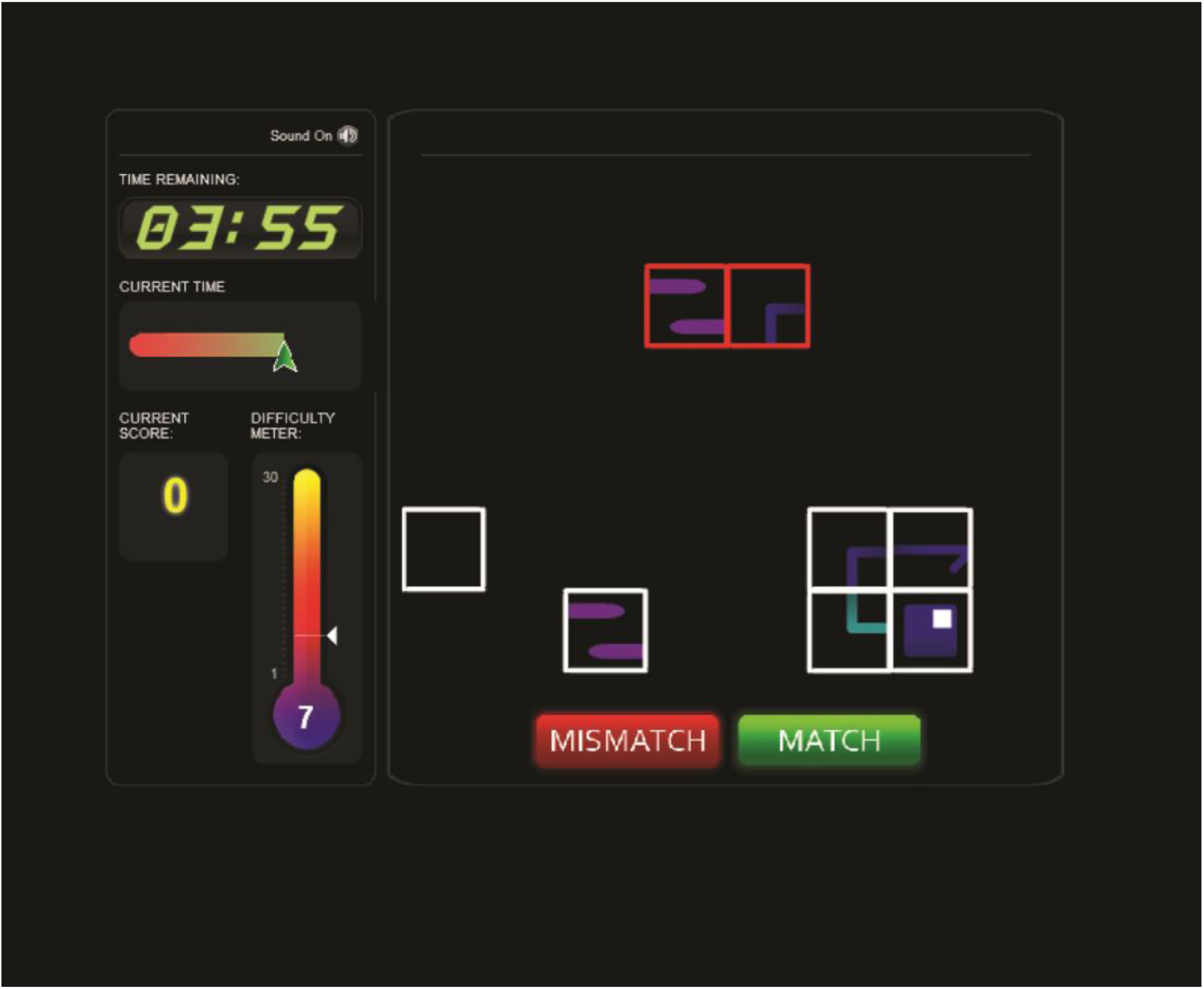
An example screen from the jigsaw task. Participants decided whether the red jigsaw at the top of the screen could be made from the pieces below. This example shows a ‘match’.

**S4 Figure.**
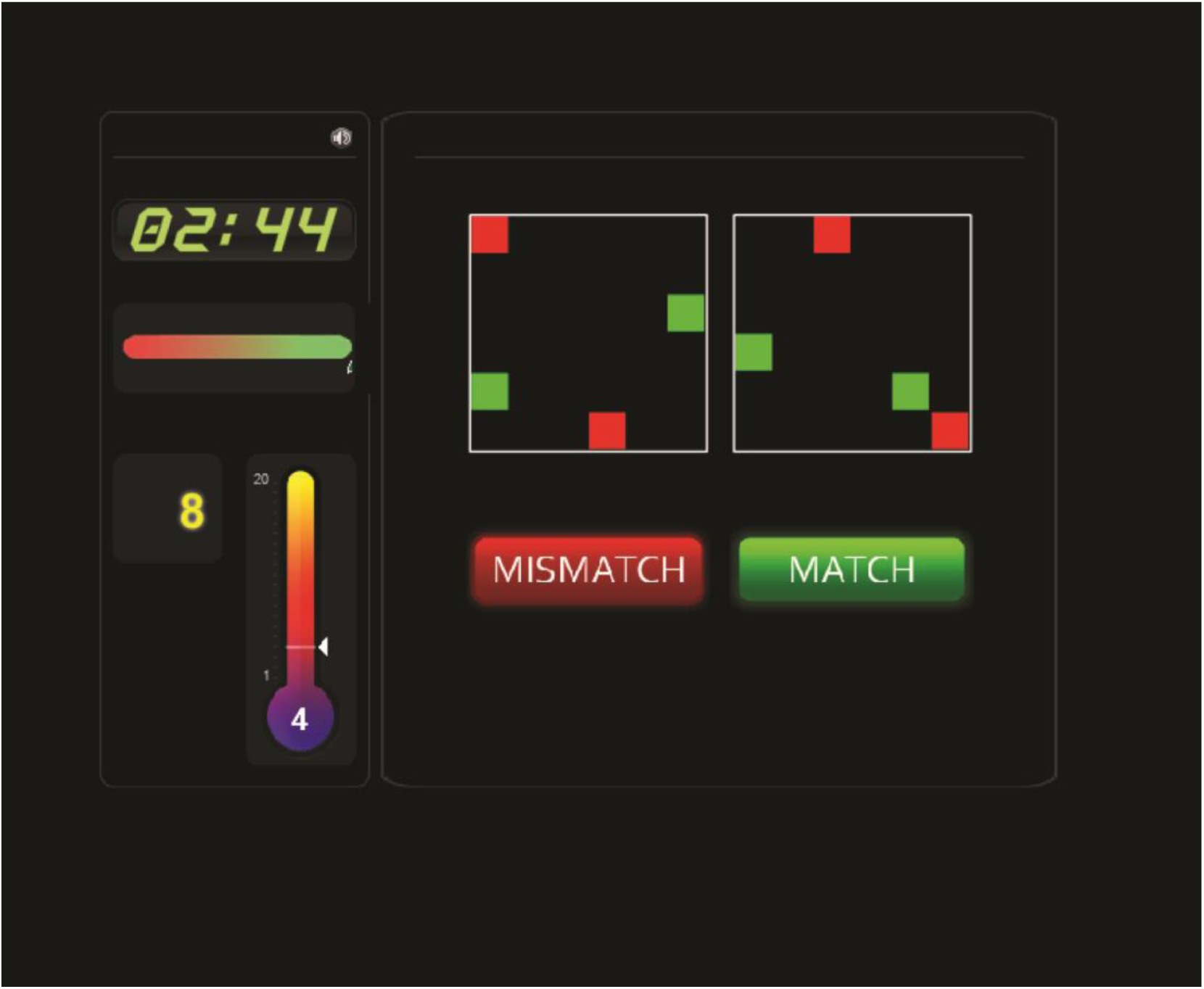
An example of the rotations task. This shows a ‘mismatch’ trial as if the right sided box was rotated so that the red boxes aligned the green boxes would not align with those in the left hand box.

**S5 Figure.**
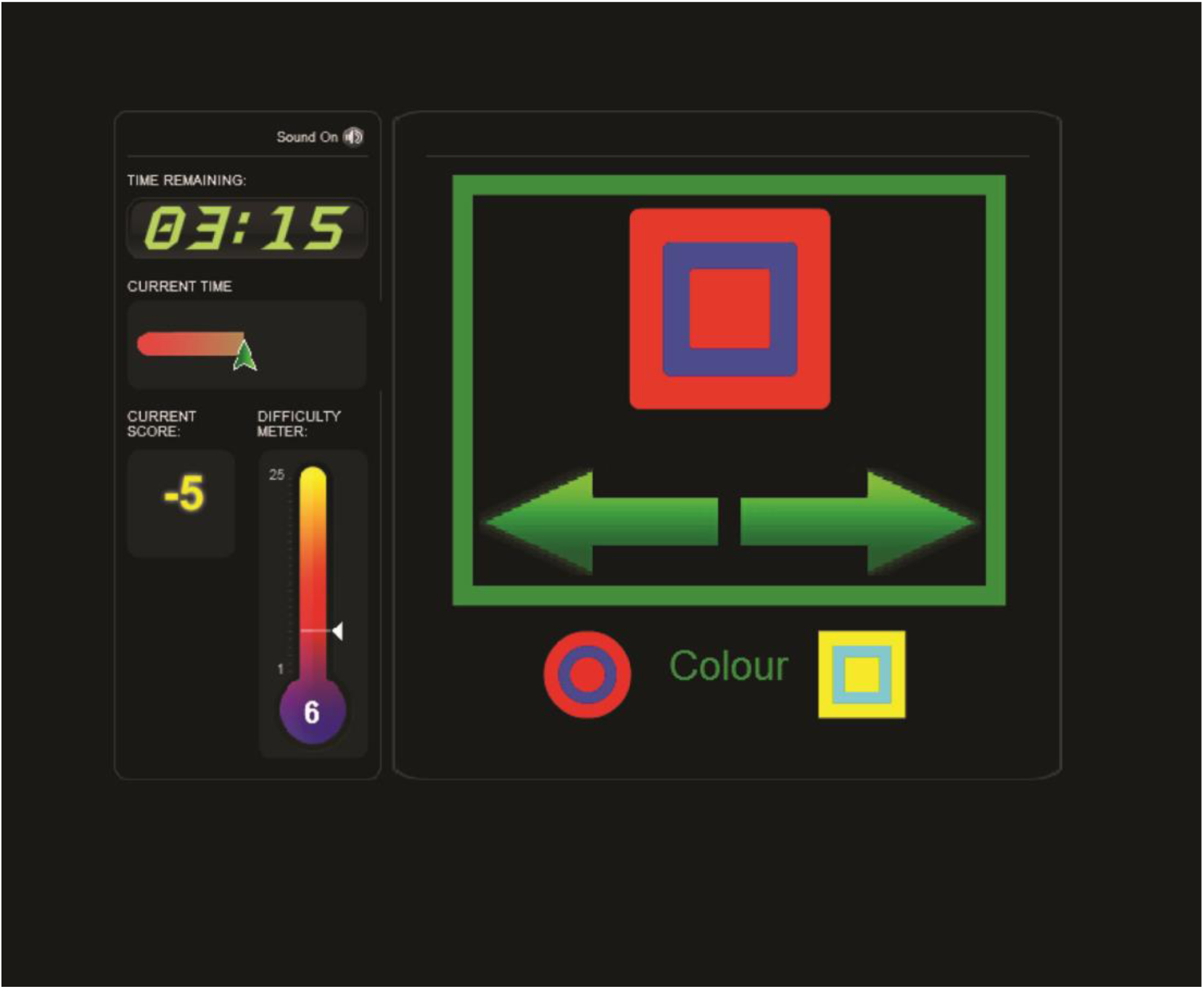
An example of the button sorting task. Here the participant must sort the top stimulus by colour, the correct response would be to click on the left pointing arrow.

